# Assessing the analytical validity of SNP-chips for detecting very rare pathogenic variants: implications for direct-to-consumer genetic testing

**DOI:** 10.1101/696799

**Authors:** Michael N Weedon, Leigh Jackson, James W Harrison, Kate S Ruth, Jessica Tyrrell, Andrew T Hattersley, Caroline F Wright

## Abstract

**Objectives:** To determine the analytical validity of SNP-chips for genotyping very rare genetic variants.

**Design:** Retrospective study using data from two publicly available resources, the UK Biobank and the Personal Genome Project.

**Setting:** Research biobanks and direct-to-consumer genetic testing in the UK and USA.

**Participants:** 49,908 individuals recruited to UK Biobank, and 21 individuals who purchased consumer genetic tests and shared their data online via the Personal Genomes Project.

**Main outcome measures:** We assessed the analytical validity of genotypes from SNP-chips (index test) with sequencing data (reference standard). We evaluated the genotyping accuracy of the SNP-chips and split the results by variant frequency. We went on to select rare pathogenic variants in the *BRCA1* and *BRCA2* genes as an exemplar for detailed analysis of clinically-actionable variants in UK Biobank, and assessed BRCA-related cancers (breast, ovarian, prostate and pancreatic) in participants using cancer registry data.

**Results:** SNP-chip genotype accuracy is high overall; sensitivity, specificity and precision are all >99% for 108,574 common variants directly genotyped by the UK Biobank SNP-chips. However, the likelihood of a true positive result reduces dramatically with decreasing variant frequency; for variants with a frequency <0.001% in UK Biobank the precision is very low and only 16% of 4,711 variants from the SNP-chips confirm with sequencing data. Results are similar for SNP-chip data from the Personal Genomes Project, and 20/21 individuals have at least one rare pathogenic variant that has been incorrectly genotyped. For pathogenic variants in the *BRCA1* and *BRCA2* genes, the overall performance metrics of the SNP-chips in UK Biobank are sensitivity 34.6%, specificity 98.3% and precision 4.2%. Rates of BRCA-related cancers in individuals in UK Biobank with a positive SNP-chip result are similar to age-matched controls (OR 1.28, P=0.07, 95% CI: 0.98 to 1.67), while sequence-positive individuals have a significantly increased risk (OR 3.73, P=3.5×10^−12^, 95% CI: 2.57 to 5.40).

**Conclusion:** SNP-chips are extremely unreliable for genotyping very rare pathogenic variants and should not be used to guide health decisions without validation.

**SUMMARY BOX:** *Section 1: What is already known on this topic:* SNP-chips are an accurate and affordable method for genotyping common genetic variants across the genome. They are often used by direct-to-consumer (DTC) genetic testing companies and research studies, but there several case reports suggesting they perform poorly for genotyping rare genetic variants when compared with sequencing.

*Section 2: What this study adds:* Our study confirms that SNP-chips are highly inaccurate for genotyping rare, clinically-actionable variants. Using large-scale SNP-chip and sequencing data from UK Biobank, we show that SNP-chips have a very low precision of <16% for detecting very rare variants (i.e. the majority of variants with population frequency of <0.001% are false positives). We observed a similar performance in a small sample of raw SNP-chip data from DTC genetic tests. Very rare variants assayed using SNP-chips should not be used to guide health decisions without validation.

## INTRODUCTION

Single gene disorders are usually caused by genetic variants that are very rare in the population (<1 in 10,000 individuals) [1]. Finding one of these rare pathogenic variants confers a very high probability of disease in an individual and their family that requires referral for clinical follow-up. For example, a confirmed pathogenic variant in one of the breast-cancer genes, *BRCA1* or *BRCA2,* would need urgent follow-up with additional screening and potentially prophylactic surgical mastectomy and oophorectomy [2]. Molecular diagnostic laboratories typically use highly accurate DNA sequencing technologies to test for these types of rare pathogenic variants [3,4].

SNP-chips are DNA microarrays that test genetic variation at many hundreds-of-thousands of specific locations across the genome [5]. They were initially designed for testing single nucleotide polymorphisms (SNPs) that are common in the population (> 1 in 100 individuals). SNP-chips have proven to be excellent for studying common genetic variation, which can be used to assess ancestry [6] as well as predisposition to many complex multifactorial diseases such as Type 2 diabetes [7,8]. Amongst the genetics community, it is generally well recognised that SNP-chips perform poorly for genotyping rare genetic variants [9–11] due to their reliance upon data clustering (Figure 1). Clustering data from multiple individuals with similar genotypes works very well when variants are common, as there are large numbers of datapoints to cluster (Figure 1a). But clustering becomes harder as the number of individuals with a particular genotype decreases; most individuals will have the reference genotype, and it is extremely difficult to distinguish a true variant from experimental noise where there is only a single carrier present (Figure 1b).

**Figure 1.**
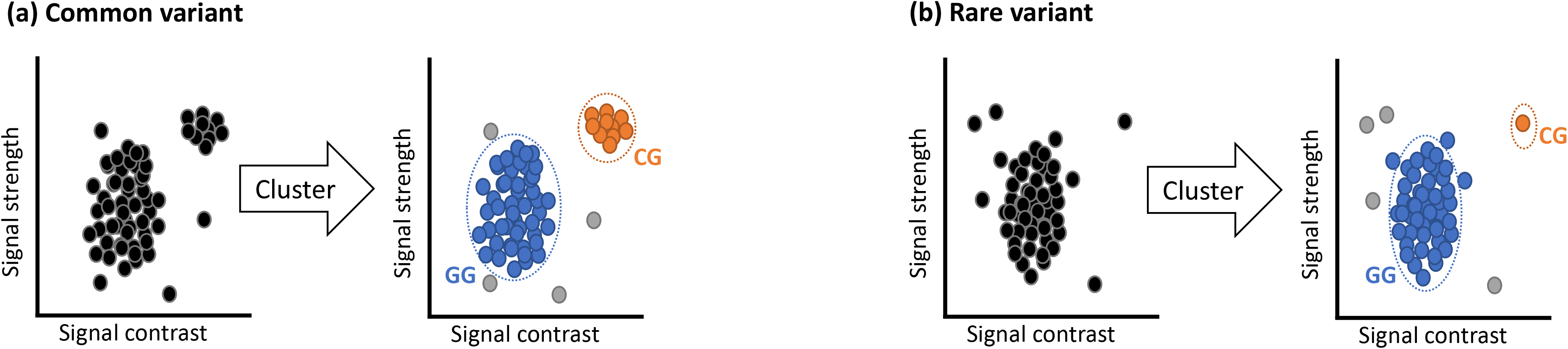
Explanation of SNP-chips genotyping technology. Example cluster plots for **(a)** a common variant and **(b)** a rare variant. Each circle represents one person’s DNA assayed at a specific position on the SNP-chip where there is a known variant (G to C). Automated clustering across multiple individuals is used to determine which DNA base is present in each person at that position. Blue circles in the main cluster represent the most common reference base (G), orange circles represent the heterozygous variant (C), and grey circles represent uncertain or missing results due to experimental noise.

Despite this problem, in recent years many SNP-chip designs, including those used by some DTC companies, have been augmented to include rare pathogenic variants that cause single gene disorders. As a consequence, consumers of DTC tests are increasingly being screened for numerous rare single gene disorders and are potentially receiving medically-actionable results, which they often take to healthcare professionals for advice [12] (Figure 2). False positive results for rare clinically-actionable variants detected by DTC SNP-chips have been described in practice guidance [13], several case reports [14,15] and two small case series [16,17]. However, there is no published systematic evaluation of the accuracy of SNP-chips for assaying rare genetic variants. It has been estimated that >26 million individuals had accessed DTC genetic testing at the start of 2019 [18], so it is crucial to know how accurate these results are likely to be in order to interpret rare pathogenic variants detected using SNP-chips.

**Figure 2.**
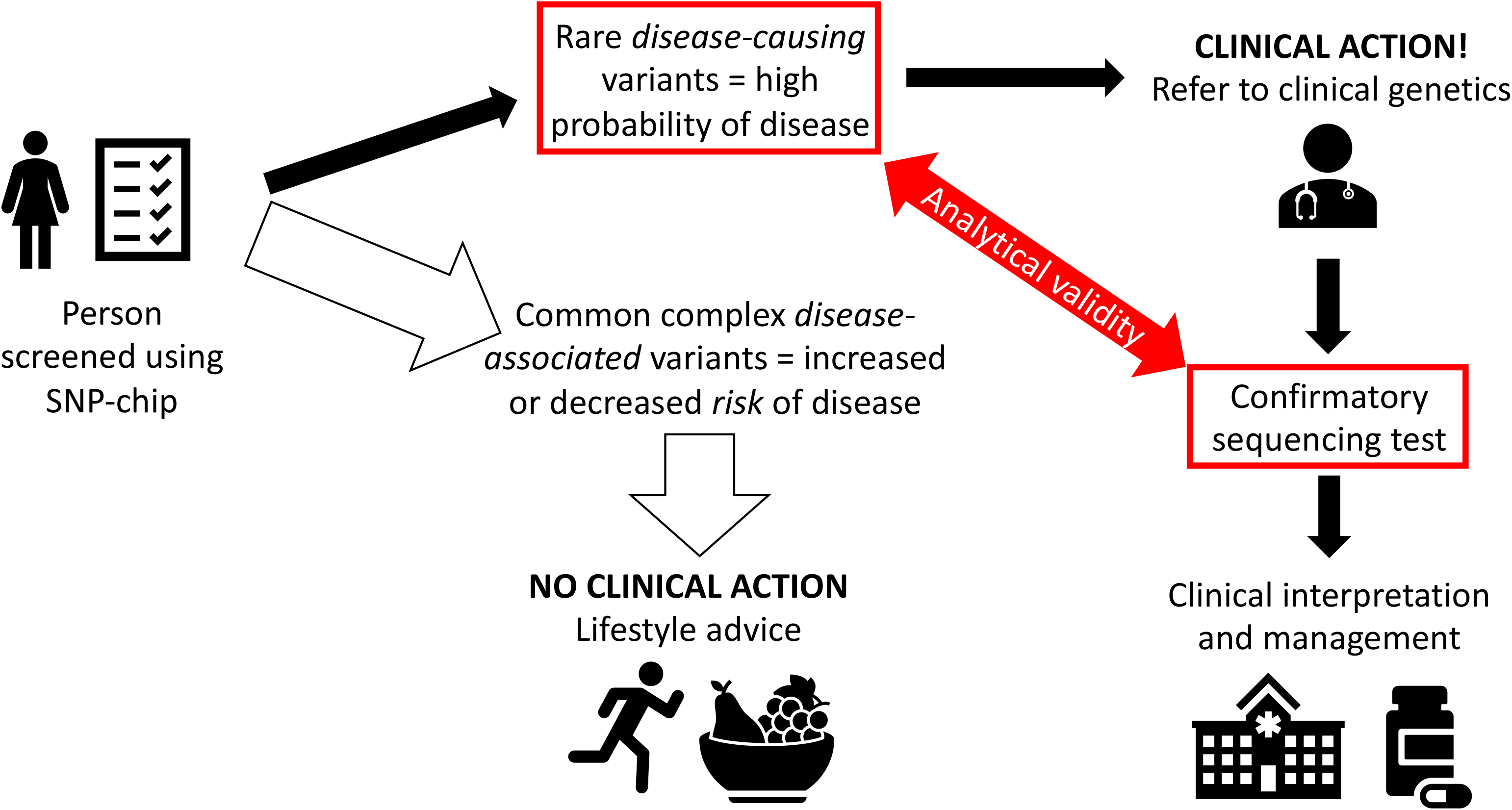
Current medical context of SNP-chip screening.

Here, we use sequencing data from 49,908 UK Biobank (UKB) [10] participants as a reference standard to do a large-scale, systematic evaluation of how well SNP-chips detect rare genetic variants. We use rare pathogenic variants in the *BRCA1* and *BRCA2* genes (collectively termed henceforth as BRCA) that cause hereditary cancers as an exemplar to evaluate the performance of SNP-chips in UKB for genotyping clinically-actionable variants. We replicate our findings using data from 21 individuals who have had DTC genetic testing and shared their data online via the Personal Genomes Project (PGP) [19]. We show that SNP-chips are extremely unreliable for genotyping very rare pathogenic variants.

## METHODS

### Study design

We performed a retrospective analytical validity study of SNP-chips (index test) using next generation sequencing (reference standard) from UKB and PGP participants in whom both datasets were available.

### Participants

We studied 49,960 individuals (55% female) from UK Biobank. The UKB is a population-based research cohort of ~500,000 participants recruited in the UK between 2006 and 2010. Approximately 9.2 million individuals aged 40-69 years who lived within 40 km of one of 22 assessment centres were invited and 5.5% participated [10]. The PGP is a community project where participants are invited to publicly share their genetic data [19].

### Test methods

We compared variants genotyped using SNP-chips (index test) with next generation sequencing data (reference standard) in the same individual.

### UKB

Next generation sequencing data were available on 49,960 individuals, of whom 49,908 also had QC-passed SNP-chip data. SNP-chip data were generated centrally by UKB and the exome sequencing data were generated externally by Regeneron and are returned to the UKB resource as part of an external access application request [20]. A subset of 4,037 individuals were previously genotyped using the Applied Biosystems UK BiLEVE Axiom Array by Affymetrix (807,411 genetic markers), and 45,871 individuals were previously genotyped using the Applied Biosystems UK Biobank Axiom^®^ Array (825,927 genetic markers) that shares 95% of its marker content with the BiLEVE [10]. Individuals were genotyped in 106 batches of ~5000 samples. We included samples that passed central UKB quality control (QC) on either of the UKB SNP-chips and used standard quality metrics to exclude problematic SNPs (missingness rate <5% and Hardy Weinberg *P*<1×10^−6^) [11]. We used the UCSC genome browser liftover tool to convert SNP-chip variant positions that were reported in hg37 to hg38 coordinates for direct comparison with sequencing data.

### PGP

We analysed publicly available datasets within the PGP (https://my.pgp-hms.org/) to determine which individuals had both DTC SNP-chip and sequencing data in hg37. We subsequently downloaded SNP-chip data (provided by 23andMe^®^ from 2012-2019 using Illumina arrays) and genome sequencing data (provided by Veritas Genetics) for 21 individuals.

### Analyses

We compared variants directly genotyped on the SNP-chips with the equivalent positions in the sequencing data from the same individual, excluding sites that were not well sequenced. For genome-wide comparison with SNP-chip genotypes, we only included directly genotyped single nucleotide variants with genomic positions present in the gVCF files and covered by >15 reads in the sequencing data. For UKB data, we used the minor allele frequency (MAF) from all 488,377 SNP-chip genotyped individuals in UKB [10]. For PGP data, we used the MAFs from gnomAD [21] and the 1000 genomes project [22]. For common and rare variant subsets in UKB, we tested the genotyping quality of heterozygous variants on the SNP-chips versus sequencing and calculated average performance metrics per variant for each subset. Representative 2×2 tables for a common, rare and very rare variant are shown in Supplementary Figure 1.

For detailed gene-specific comparison with SNP-chip genotypes in UKB, we included directly genotyped single nucleotide variants, as well as small insertions and deletions in the *BRCA1* and *BRCA2* genes. Variants were defined as pathogenic if they were either predicted to result in a truncated protein or had previously been classified as likely or definitely pathogenic in the ClinVar database [23]; variants with conflicting reports of pathogenicity that included pathogenic assertions made within the last 5 years were included. Sequencing data was visually examined using the Integrative Genomics Viewer (IGV) [24] to determine whether the variant was present or not. Cancer registry data for breast, ovarian, prostate and pancreatic cancer was extracted for all participants. Logistic regressions were carried out to assess the relationship between test-positive participants and any BRCA-related cancer.

Results are presented in accordance with the standard framework for the validation and verification of clinical molecular genetic tests [25] and STARD guidelines for reporting diagnostic accuracy studies [26], using sensitivity, specificity and precision to evaluate assay performance.

### Patient and public involvement

No patients or the public were directly involved in the design or implementation of this study as we used data previously generated. As part of the consent process for UKB, NHS patients gave their consent for the collection, storage and use of their genetic data and medical records by all approved researchers. All of the participant records are linked-anonymised. We will disseminate this research through the UK Biobank network. The PGP participants have freely shared their data online to be used by the research community.

## RESULTS

### Performance of SNP-chips appears excellent when assessed for all variants

In the 49,908 UK Biobank individuals, we compared genotypes for 108,574 single nucleotide variants that were classified as heterozygous by the SNP-chip to reference standard sequencing data. Of the 49,908 individuals, 45,871 were genotyped using the Axiom^®^ chip and 4,037 using the BiLEVE chip. Overall performance across both chips for all variants is very good (Table 1), with 3.1 ×10^8^ true positives, 4.6×10^9^ true negatives, 3.2×10^6^ false positives and 2.7×10^6^ false negatives. Performance of both chips for genotyping common SNPs with a frequency of >1% is especially good (Table 1): Axiom^®^ chip (average sensitivity 99.8%, specificity 99.7% and precision 99.0%) and BiLEVE chip (average sensitivity 99.7%, specificity 99.7% and precision 98.7%).

**Table 1.** Performance of UKB SNP-chips versus sequencing. Results are split by SNP-chip and variant group: all variants, common variants (MAF >1%), very rare variants (MAF <0.001%), and pathogenic BRCA variants. 95% confidence intervals are given in brackets. Insertions and deletions are excluded except in the BRCA analysis. (MAF = Minor Allele Frequency)

| SNP-chip | Individuals | Dataset | Sensitivity (%) | Specificity (%) | Precision (%) |
| --- | --- | --- | --- | --- | --- |
| UKB Axiom® | 45,871 | All variants | 96.3 (96.2 to 96.4) | 99.9 [99.9 to 99.9] | 85.9 [85.7 to 86.1] |
|  |  | MAF >1% | 99.8 [99.8 to 99.8] | 99.7 [99.7 to 99.8] | 99.0 [98.9 to 99.1] |
|  |  | MAF <0.001% | 29.5 [27.4 to 31.5] | 99.9 [99.9 to 99.9] | 16.1 [14.9 to 17.3] |
|  |  | BRCA pathogenic | 33.0 [22.8 to 43.3] | 99.7 [99.6 to 99.7] | 16.9 [12.9 to 22.0] |
| UKB BiLEVE | 4,037 | All variants | 96.9 [96.8 to 97.0] | 99.9 [99.9 to 99.9] | 91.1 [91 to 91.2] |
|  |  | MAF >1% | 99.7 [99.7 to 99.8] | 99.7 [99.7 to 99.8] | 98.7 [98.6 to 98.8] |
|  |  | MAF <0.001% | 4.4 [3.1 to 5.8] | 99.9 [99.9 to 99.9] | 9.4 [6.8 to 12.0] |
|  |  | BRCA pathogenic | 50.0 [18.7 to 81.3] | 82.7 [81.5 to 83.9] | 0.7 [0.4 to 1.3] |

### False positive results from SNP-chips greatly increases with decreasing allele frequency

The genotyping performance of the SNP-chips in UKB is strongly related to the frequency of the variant in the population (Table 1, Figure 3 and Supplementary Figure 2). There are 10,891 (Axiom^®^) and 7,408 (BiLEVE) variants on the two UKB SNP-chips with a frequency <0.001% in UKB. For these very rare variants, the sensitivity of the SNP-chips to detect heterozygous genotypes is low (29.5% for Axiom^®^ and 4.4% for BiLEVE) although, because the variant alleles are very rare, the specificity remains high (99.9% for both chips). However, the precision is strikingly reduced for rare variants compared with common variants, i.e. there is a very high rate of false positives. For the very rare variants in UKB, only 16.1% of Axiom SNP-chip heterozygous genotypes (708 true positives in 4,239 individuals at 3,422 variants) were confirmed by the sequencing data and only 9.4% (46 true positives in 518 individuals at 488 variants) for the BiLEVE chip. A similar performance was seen for very rare variants in the PGP data (Supplementary Figure 3), with a precision of 14% for variants with a population frequency <0.01% in 21 individuals (83 true positives in 21 individuals at 594 variants).

**Figure 3.**
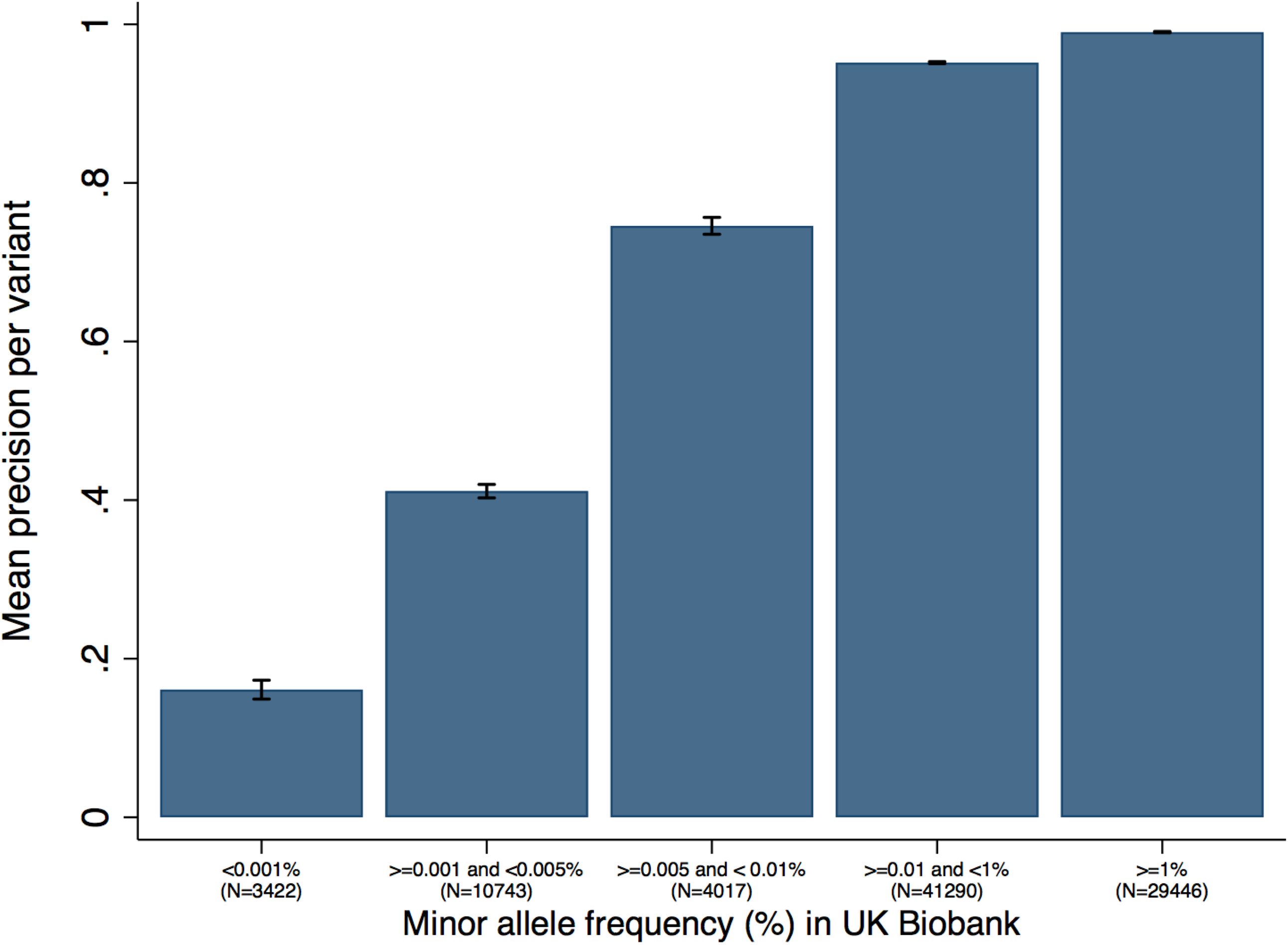
Precision of UKB Axiom^®^ SNP-chip for detecting variants at different population frequencies. A similar trend is seen with the UKB BiLEVE chip (Supplementary Figure 2) and the PGP consumer data (Supplementary Figure 3).

### Rare pathogenic variants are very poorly genotyped by SNP-chips

We went on to evaluate the performance of the SNP-chips in UKB for 1,139 pathogenic and likely pathogenic variants in BRCA that were included on the chips (Figure 4 and Supplementary Figure 4); 916 (80%) of these are rare with an allele frequency of <0.01% in UKB. The performance of both chips is very poor for genotyping pathogenic BRCA variants in UKB (overall sensitivity 34.6%, specificity 98.3% and precision 4.2%), with a very high rate of false positive variants (Table 1). Across both SNP-chips there are 425 pathogenic BRCA variants in 889 individuals in UKB. Of these, just 17 variants in 37 individuals are present in the sequencing data; the most common variant is present in 10 individuals and has conflicting and uncertain interpretations in ClinVar. A further 43 pathogenic BRCA variants are present in the sequencing data of 70 individuals but are not detected by either SNP-chip despite being assayed. The performance of both chips for genotyping pathogenic BRCA variants is very poor (Table 1): Axiom^®^ chip (sensitivity 33.0% and specificity 99.7%) and BiLEVE chip (sensitivity 50.0% and specificity 82.7%).

**Figure 4.**
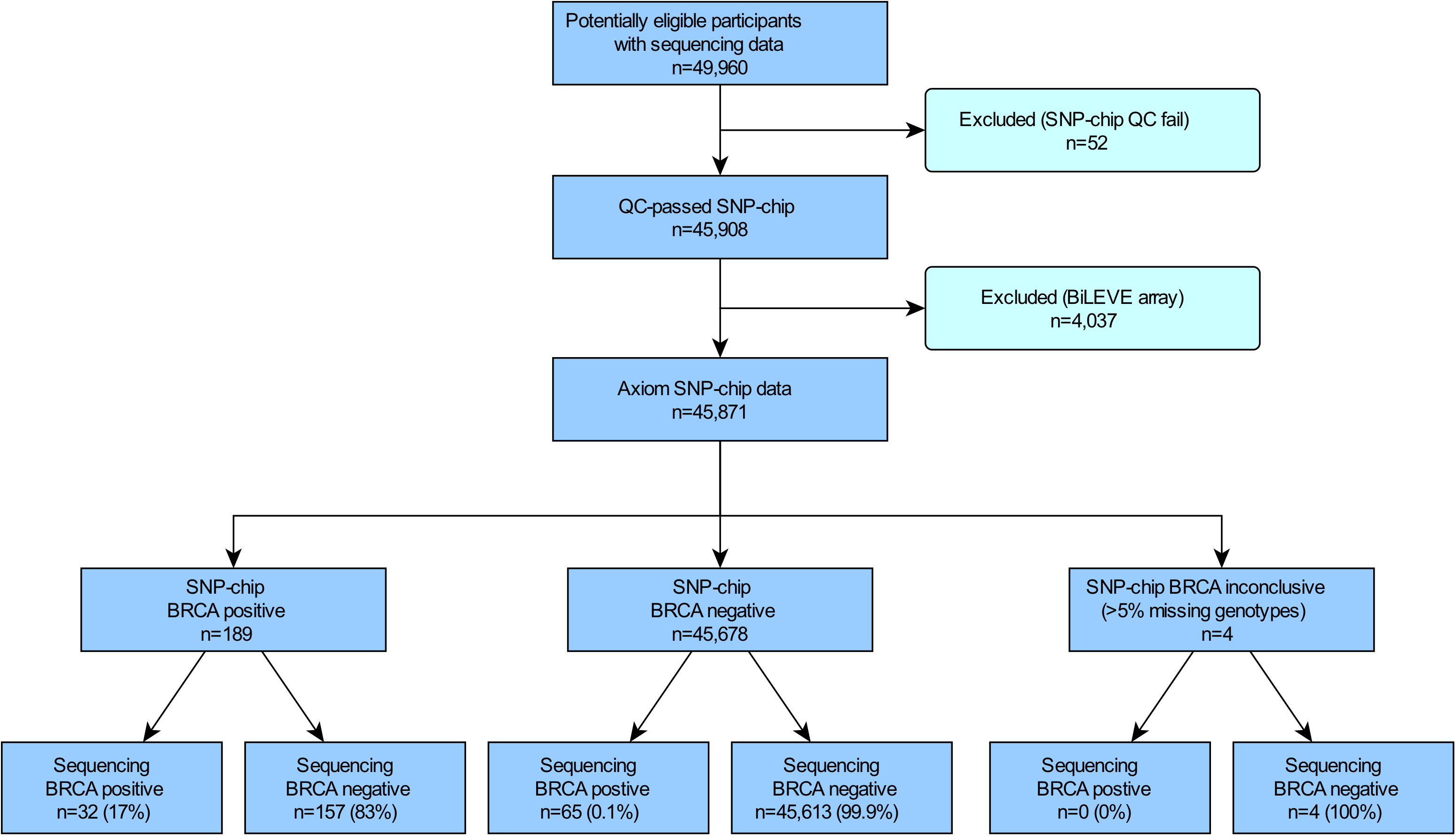
STARD diagram to report flow of participants with a pathogenic BRCA variant on the UKB Axiom^®^ chip compared with sequencing. A similar process was followed for the BiLEVE chip (Supplementary Figure 4).

We also investigated rare pathogenic/likely pathogenic variants in PGP. Across 21 individuals, 100% (47/47) of rare (MAF<0.01%) pathogenic single nucleotide variants and 74% (25/34) of likely protein-truncating insertions and deletions in known disease-causing genes are incorrectly genotyped by the SNP-chips. These include a pathogenic variant in *KCNQ2* that causes early infantile epileptic encephalopathy [27], and protein-truncating variants in *MSH2* and *MSH6* that confer a very high risk of colorectal cancer [28]. A rare pathogenic variant in *ABCC8* that causes congenital hyperinsulinism [29] is incorrectly genotyped in 43% (9/21) of individuals. Overall, 95% (20/21) of individuals investigated have a least one false positive rare pathogenic variant.

### Risk of BRCA-related hereditary cancer in individuals in UKB with a pathogenic BRCA variant genotyped by SNP-chips is similar to population risk

The risk of BRCA-related cancers in UKB individuals with a positive SNP-chip result for any pathogenic BRCA variant is similar to the age-matched risk in UKB (OR 1.28, P=0.07, 95% CI: 0.9 to 1.67). In contrast, those with a positive sequencing result – including the 107 individuals with BRCA variants assayed by either chip, plus another 137 individuals with pathogenic BRCA variants not assayed by either chip – have a markedly increased risk (OR 3.73, P=3.5×10^−12^, 95% CI: 2.57 to 5.40).

## DISCUSSION

We have shown that SNP-chips are extremely poor for correctly genotyping very rare variants compared with sequencing data. It is widely recognised that SNP-chips are not good at genotyping very rare variants [9–11] so some recent SNP-chips were designed only to assay low and intermediate frequency coding variants (>1 in 5000), for which the SNP-chips perform relatively well. However, increasingly SNP-chips are being augmented with very rare pathogenic variants which, as we show, are not well genotyped. This low analytical validity for rare genetic variants is an inherent issue with the clustering method upon which SNP-chips rely (Figure 1). Genotyping batch size will therefore also affect accuracy, with fewer individuals per batch leading to more genotyping errors for rare variants. As a result, although the performance of chips from different manufacturers may differ somewhat due to different underlying chemistries [5], our findings are likely to be generalisable to any SNP-chip, and indeed the results are similar across different SNP-chips used in both our UKB and PGP datasets. Sequencing data are not affected by the same technical issues as SNP-chips, and thus provide a much more accurate method for genotyping rare variants.

The poor genotyping quality of SNP-chips can be remedied through improved probe design, custom variant detection definitions [30], using multiple probes for individual variants or using positive laboratory controls to improve variant clustering. Many DTC companies use these additional quality control methods, supplemented by validation of important variants using DNA sequencing, to improve the accuracy of variants they advertise and report directly to consumers. However, most DTC companies (including those focused on ancestry or other non-medical traits) also allow customers to download and analyse their raw data, which will often include many thousands of additional rare variants assayed on the SNP-chip that have not undergone stringent quality control and are therefore much more likely to be false positives [15]. A recent study found that 89% of consumers of DTC genetic tests download their raw data and 94% of those use at least one third-party interpretation service to analyse the results [31]. We have therefore focused on the DTC scenario because errors in raw SNP-chip data could cause significant harm to patients without further validation. However, erroneous results from SNP-chips used in research biobanks can also lead to false associations [32] and wasted resource in the development of new treatments against the wrong targets [33].

Using a large population research cohort and a small consumer genetic testing cohort, we have shown that positive results from SNP-chips for very rare variants are more likely to be wrong than right. For pathogenic BRCA variants in UKB, the SNP-chips have an extremely low precision (1-17%) when compared with sequencing. Were these results to be fed-back to individuals, the clinical implications would be profound. Women with a positive BRCA result face a lifetime of additional screening and potentially prophylactic surgery that is unwarranted in the case of a false positive result. Conversely, although the false negative rate of SNP-chips is generally low, many very rare pathogenic variants are not included in the design and will therefore be missed [34]. Women who receive false negative BRCA result but have a strong family history of breast and/or ovarian cancer are at extremely high risk of developing cancer that could be greatly reduced through preventive surgeries and other interventions [35].

We urge clinicians to validate any SNP-chip results from DTC companies or research biobanks using a standard diagnostic test prior to recommending any clinical action. In addition, individuals with symptoms or a family history of breast and/or ovarian cancer who have received a negative SNP-chip result should not be reassured that their risk is low [13] and standard referral guidelines should be followed for diagnostic testing (see https://cks.nice.org.uk/breast-cancer-managing-fh). We suggest that, for variants that are very rare in the population being tested, genotyping results from SNP-chips should not be routinely reported back to individuals or used in research without validation. Clinicians and researchers should be aware of the poor performance of SNP-chips for genotyping very rare genetic variants to avoid counselling patients inappropriately or investing limited resources into investigating false associations with badly genotyped variants.

## Supporting information

Definitions Box

## ACKNOWLEDGEMENTS

The authors wish to thank Dr Charles Warden for pointing us towards the PGP resource and Dr Tim McDonald for useful advice on clinical performance metrics. We also wish to thank participants of both UK Biobank and the Personal Genomes Project for generously donating their data for research. This research has been conducted using the UK Biobank Resource under Application Number 49847 and 871, using the University of Exeter High-Performance Computing (HPC) facility.

## CONTRIBUTORS

CFW, MNW and ATH designed the study. CFW wrote the first draft of the manuscript. All authors reviewed and edited the manuscript. MNW, LJ, JWH, KSR, JT, ATH and CFW were involved in data processing, statistical analysis, and interpretation. CFW is the guarantor.

## ETHICAL APPROVAL

UK Biobank has received ethics approval from the National Health Service National Research Ethics Service (ref 11/NW/0382).

## TRANSPARENCY STATEMENT

The lead author (the manuscript’s guarantor) affirms that this manuscript is an honest, accurate, and transparent account of the study being reported; that no important aspects of the study have been omitted; and that any discrepancies from the study as planned (and, if relevant, registered) have been explained.

## DATA SHARING

The data reported in this paper are available via application directly to the UK Biobank.

This is an Open Access article distributed in accordance with the terms of the Creative Commons Attribution (CC BY 3.0) license, which permits others to distribute, remix, adapt and build upon this work, for commercial use, provided the original work is properly cited. See: http://creativecommons.org/licenses/by/3.0/.

