## Supplementary Figures for "Assessing the analytical validity of SNP-chips for detecting very rare pathogenic variants: implications for direct-to-consumer genetic testing"

**Supplementary Figure 1.** Representative 2x2 tables for a common, rare and very variant in UKB.

**REFERENCE STANDARD (SEQUENCING)**

| INDEX TEST (SNP-CHIP) | **Common variant (UKB MAF = 11%)** | | **Rare variant (UKB MAF = 0.1%)** | | **Very rare variant (UKB MAF = 0.0007%)** | |
|---|---|---|---|---|---|---|
| | Variant positive | Variant negative | Variant positive | Variant negative | Variant positive | Variant negative |
| Variant positive | 9109 | 97 | 130 | 1 | 0 | 1 |
| Variant negative | 18 | 36505 | 0 | 45718 | 0 | 45870 |

**Supplementary Figure 2.** Precision of UKB BiLEVE SNP-chip for detecting variants at different population frequencies.

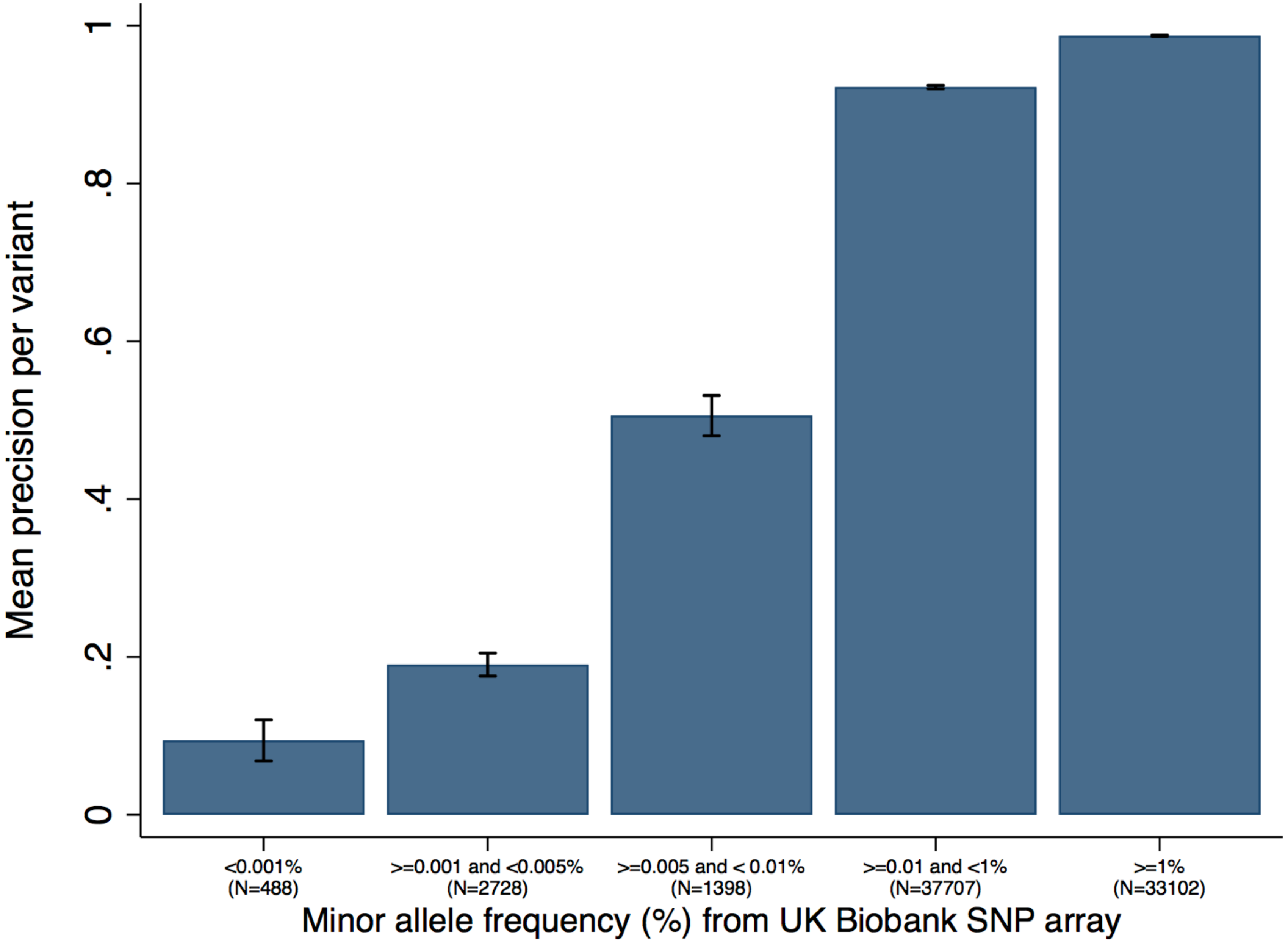

**Supplementary Figure 3.** Precision of PGP SNP-chips for detecting variants at different population frequencies.

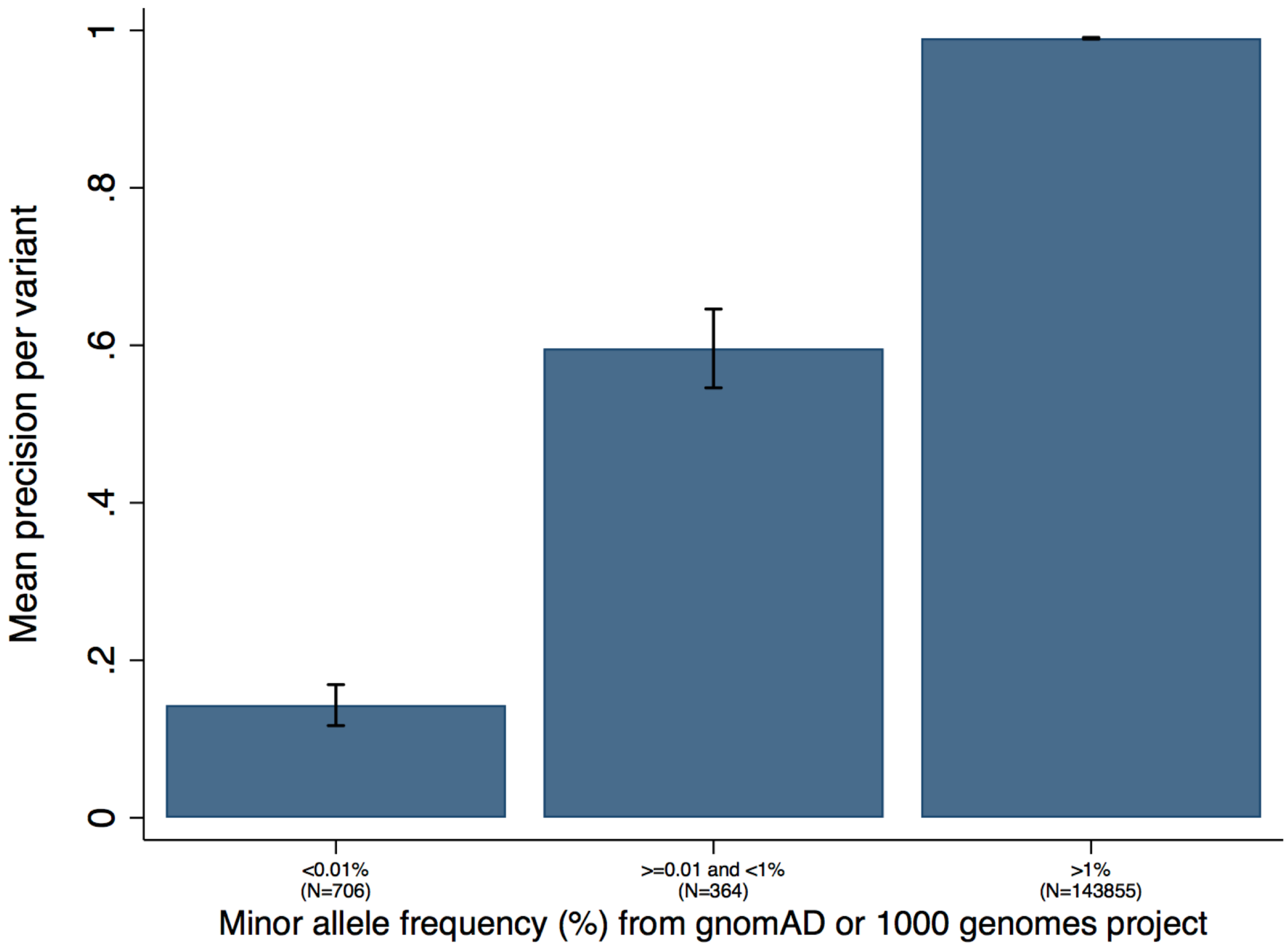

**Supplementary Figure 4.** STARD diagram to report flow of participants with a pathogenic BRCA variant on the UKB BiLEVE® chip compared with sequencing.

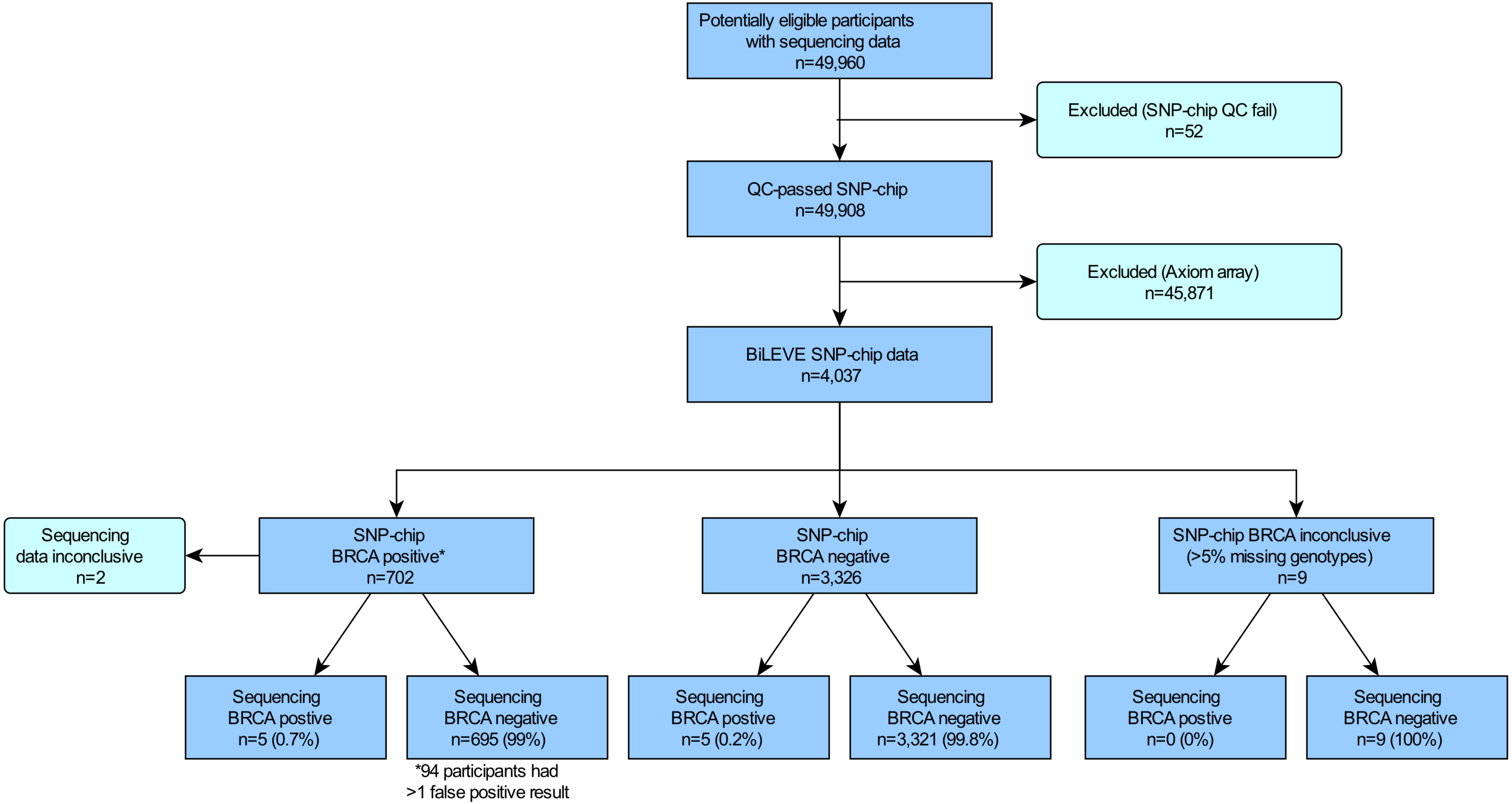
