## Supplementary material for "Assessing the analytical validity of SNP-chips for detecting very rare pathogenic variants: implications for direct-to-consumer genetic testing": Definitions Box

- **Single gene disorders** = diseases caused by, or with a high probability of developing due to, a rare genetic variant in a specific single gene.
- **Allele** = each of two or more alternative forms of DNA that are found at the same location on a chromosome.
- **Genome** = the complete set of genes or genetic material present in a cell or organism.
- **Variant** = position in the genome where an individual differs from the reference human genome by a single base pair change, i.e. a substitution of a single letter of DNA. A variant may be rare or common in the population.
- **SNP (Single Nucleotide Polymorphism)** = type of single nucleotide variant that is common and present in greater than 1% of the population (pronounced "snip").
- **SNP-chip** = DNA microarray that is used to genotype known genetic variants (typically SNPs) in the population.
- **Genotyping** = method for determining the base pair (A, G, T or C) present at a specific location in a person's DNA. This be achieved by various methods including SNP-chips or sequencing.
- **DNA sequencing** = method for determining the order of base pairs in a DNA sample and the variants within it.
- **Sensitivity** = proportion of variants detected by the reference test that are also found by the index test [TP/(TP+FN)].
- **Specificity** = proportion of variants not detected by the reference test that are also found to be normal by the index test [TN/(FP+TN)].
- **Precision =** proportion of variants found by the index test that are confirmed by the reference test [TP/(TP+FP)].
